# Time Cells in the Retrosplenial Cortex

**DOI:** 10.1101/2024.03.01.583039

**Authors:** Dev Laxman Subramanian, David M. Smith

## Abstract

The retrosplenial cortex (RSC) is a key component of the brain’s memory systems, with anatomical connections to the hippocampus, anterior thalamus, and entorhinal cortex. This circuit has been implicated in episodic memory and many of these structures have been shown to encode temporal information, which is critical for episodic memory. For example, hippocampal time cells reliably fire during specific segments of time during a delay period. Although RSC lesions are known to disrupt temporal memory, time cells have not been observed there. In the present study, we examined the firing patterns of RSC neurons during the intertrial delay period of two behavioral tasks, a blocked alternation task and a cued T-maze task. For the blocked alternation task, rats were required to approach the east or west arm of a plus maze for reward during different blocks of trials. Because the reward locations were not cued, the rat had to remember the goal location for each trial. In the cued T-maze task, the reward location was explicitly cued with a light and the rats simply had to approach the light for reward, so there was no requirement to hold a memory during the intertrial delay. Time cells were prevalent in the blocked alternation task, and most time cells clearly differentiated the east and west trials. We also found that RSC neurons could exhibit off-response time fields, periods of reliably inhibited firing. Time cells were also observed in the cued T-maze, but they were less prevalent and they did not differentiate left and right trials as well as in the blocked alternation task, suggesting that RSC time cells are sensitive to the memory demands of the task. These results suggest that temporal coding is a prominent feature of RSC firing patterns, consistent with an RSC role in episodic memory.

## Introduction

The retrosplenial cortex (RSC) is a key component of the brain’s memory system, with anatomical connections to the hippocampus, anterior thalamus, and entorhinal cortex (Sugar et al., 2011; Van Groen et al., 1993; Van Groen & Wyss, 2003; Wyss & Van Groen, 1992). This circuit has been implicated in episodic memory (Aggleton et al., 2023) and many of these structures have been shown to encode temporal information, which is critical for episodic memory. For example, time cells, which reliably fire during a specific epoch of time during a delay period in a manner analogous to the way that place cells fire in a specific part of the environment, have been found in CA1 (Gill et al., 2011; Kraus et al., 2013; MacDonald et al., 2011; Mau et al., 2018; Pastalkova et al., 2008), CA3 (Salz et al., 2016), the medial entorhinal cortex (Heys & Dombeck, 2018; Kraus et al., 2015), the prefrontal cortex (Ning et al., 2022; Tiganj et al., 2017), and the striatum (Akhlaghpour et al., 2016; Mello et al., 2015).

Time cells have not been reported in the RSC, although there is evidence that the RSC is involved in temporal memory. Amnesic patient T.R., who had an RSC lesion, was found to have striking deficit in temporal memory which was not attributable to generally poor memory for non-temporal information (Bowers et al., 1988). Gabriel and colleagues (Freeman et al., 1996; Smith et al., 2004) proposed a theoretical account of how learning– and time-dependent changes in RSC firing patterns could encode the spatio-temporal context of a learning situation (for review see Smith et al., 2018). RSC neurons have also been found to simulate future goal locations in rats (Miller et al., 2019). There is also circumstantial evidence in the form of sequence coding in the RSC, where neurons have been found to selectively encode specific sequences of responses and movements through space (Alexander & Nitz, 2015, 2017) and these authors predicted the likely occurrence of time cells in the RSC (Alexander et al., 2020).

In the present study, we re-examined archival neuronal firing data from the RSC recorded during two different behavioral tasks. Both tasks involve intertrial delay periods where time cells might be found, but they have differing memory requirements. One of these tasks involves spatial alternation across blocks of trials on a plus maze (Smith & Mizumori, 2006). In this task, rats must remember the reward location for the upcoming trial, and hippocampal neurons were previously found to exhibit time cell firing in this task (Gill et al., 2011). The second task is a beacon navigation task in which a salient light cue signaled the location of the reward on a T-maze and the rats simply had to approach the light for reward (Vedder et al., 2017).

## Methods

### Subjects and Surgical Procedures

The subjects were 15 adult male Long-Evans rats with 10 of them used in the blocked alternation task and 5 of them used in the cued T-maze task. For this study, we analyzed data from previous studies which focused on spatial firing patterns of RSC neurons during the trials (Smith & Mizumori, 2006; Vedder et al., 2017). Here, we focus our analysis on the firing patterns during the intertrial delay period. All rats underwent stereotaxic surgery to implant recording electrodes targeting the granular b subregion of the RSC (Rgb), also referred to as the posterior cingulate cortex or Brodmann’s Area 29c (3.0–6.0 mm posterior to bregma, 0.5mm lateral and at least 1.0 mm ventral to the dorsal surface, Paxinos & Watson, 1998). Rats were given at least one week to recover from surgery before beginning training. All procedures complied with guidelines established by the Cornell University Animal Care and Use Committee.

### Behavioral procedure and neuronal recording

For both tasks, the maze was placed in a dimly lit room with distal visual cues and correct choices were rewarded with chocolate milk (200 ul Nesquik, Nestle Inc.). Daily recordings took place after the rats reached asymptotic performance on the tasks (blocked alternation mean = 84.07 ± 0.91%; cued T-maze mean = 94.72 ± 1.02%). For the blocked alternation task, rats were trained to retrieve rewards from the east arm of a plus maze during the first block of 15 trials and then switch to the west arm for the second block of 15 trials. This yielded 28 intertrial delay periods, 14 for each block. The start positions for each trial were randomly designated from among the 3 non-reward arms. The reward locations were not cued, so the rat was required to remember which arm was rewarded for each trial. For the cued T-maze task, rats were trained on a T-maze equipped with LED lights positioned 9 cm above each of the reward locations and the rats were trained to approach a flashing light cue for a reward. At the start of each trial, the rat was placed on the stem of the maze facing away from the choice point and one of the two light cues was turned on (flashing at 3 Hz) to indicate the reward location for that trial. The left or right reward positions were randomized across trials and importantly, this task did not have a spatial memory requirement since rats could simply approach the cued location on each trial. Each training session consisted of 40 trials, with 20 rewarded on the left and 20 on the right.

Analysis of the neuronal firing data was focused on the intertrial delay period, which began when the rat was placed on a platform (5.5 × 25 cm) adjacent to the maze after the completion of each trial and ended when the rat was picked up to be placed on the maze for the start of the next trial. The duration of the intertrial delay varied as the experimenter re-baited the maze and prepared for the next trial, but typically lasted 19–28 sec (mean = 24.14 ± 0.23 sec). The position of the platform was constant throughout training.

## Data collection and analysis

Neuronal spike data and video data were collected using the Cheetah Data Acquisition System (Neuralynx Inc., Bozeman, MT). Video data were used to establish the beginning and end of the intertrial delay period. Our dataset consisted of 383 neurons, 180 neurons recorded from 10 rats in the blocked alternation task and 203 neurons recorded from 5 rats in the cued T-maze task. We classified putative pyramidal neurons and interneurons according to spike width (see Brennan et al., 2020). This yielded 237 pyramidal neurons (mean width = 0.358 ± 0.005 msec) and 146 interneurons (mean width = 0.142 ± 0.004 msec), all of which were included in our analyses.

## Time cell classification and differential firing between conditions

We applied a Linear-Nonlinear (L-N) model (Hardcastle et al., 2017) to identify neurons that exhibited significant firing changes during discrete epochs during the intertrial delay period. Briefly, the model estimates the firing rate of an individual neuron as a function of time and determines whether the tuning is statistically significant. Specifically, we binned the spikes into 20 msec bins and assessed the degree to which firing was significantly tuned to 500 msec epochs in either condition (east or west in the blocked alternation task, right or left in the cued T-maze task). We used a two-step process to classify time cells and determine whether each time cell fired differentially in the two conditions of the task (east/west and right/left). We first classified time cells as those neurons with significant tuning in either condition. Then, for each of the resulting time cells, we identified the ‘time field’ as any epoch where the firing rate diverged from the mean by at least two standard deviations, and we ran a second L-N model to determine whether the firing during that epoch was significantly different in the east/west or left/right conditions. This approach to classifying time cells yielded similar results to methods previously applied to hippocampal neurons (Gill et al., 2011; Pastalkova et al., 2008). However, the L-N model is better suited for RSC neurons, which have high baseline firing rates and responses to task variables can consist of increased or decreased firing (e.g. Miller et al., 2019; Vedder et al., 2017), and it yielded fewer apparent misclassifications than methods optimized for hippocampal neurons.

We also computed measures of time cell quality, including a reliability measure (the proportion of trials in which the firing rate during the time field was at least 2 SD above or below the mean) and a measure of contrast reflecting the in-field vs out-of-field firing rates (absolute value of (in-out)/(in+out)). Previous studies have shown that time fields can be aligned to the beginning or end of the intertrial delay period (MacDonald et al., 2011). However, our delay periods were variable, which could obscure time fields near the end of the delay, so we also analyzed the firing data of each neuron aligned to the end of the delay period.

## Results

Many RSC neurons reliably exhibited discrete periods of elevated firing during the intertrial delay period (i.e. time fields, Fig. 1A, plots 1-8), similar to those previously reported in the hippocampus (Gill et al., 2011; Pastalkova et al., 2008). In the blocked alternation task, 36% of neurons (65/180) exhibited significant time fields in at least one of the two blocks (go east or go west). Firing during the time fields was quite reliable, with firing >2 SD above the mean during the time field on 75.4% (± 1.45) of trials, and time fields were generally brief, circumscribed segments of the delay period (mean duration = 2.95 ± 0.22 sec). These time fields were distributed throughout the intertrial delay period, but they were most prevalent during the first 5 sec of the delay (Fig. 1B). Some neurons had two time fields (n=11, e.g. Fig. 1A, plot 8) and some time fields were more closely aligned to the end of the delay period than the start (n=14, Fig. 1C).

**Figure 1.**
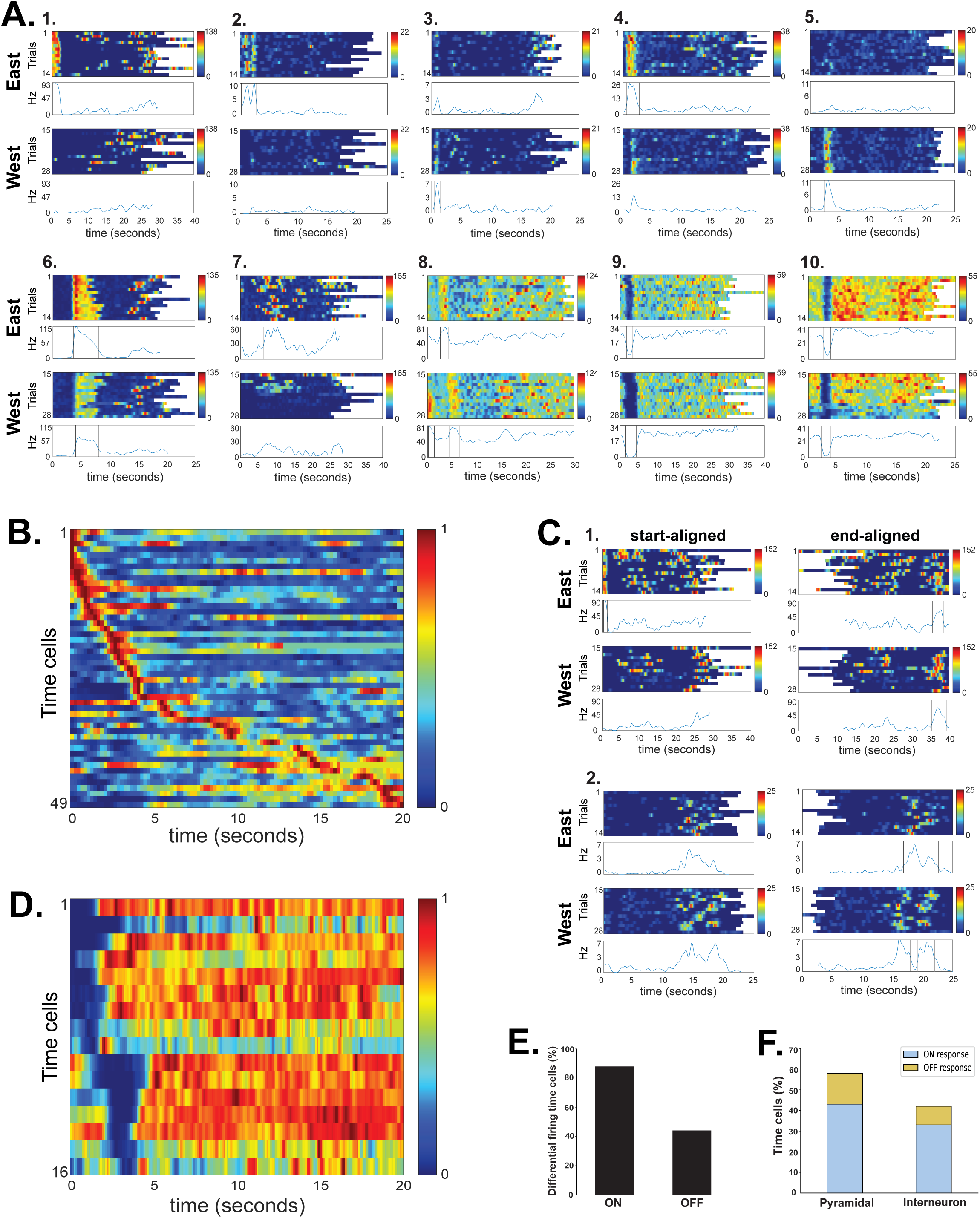
Retrosplenial Time Cells in the blocked alternation task. Plot A shows examples of RSC time cells observed in the blocked alternation task. The firing rate heatmaps illustrate trial by trial firing, shown separately for the ‘Go East’ and ‘Go West’ conditions. The line plot shows the average firing rate across trials with the two vertical black lines indicating the boundaries of the time field. Because the delay duration was variable, average firing rate was calculated only for time periods containing at least two thirds of the trials. Neurons could have more than one time field (e.g. plot 8). Plots 9 and 10 illustrate example neurons with off-response time fields, in which firing was reliably suppressed during a well-defined part of the delay. Plot B & D illustrate the temporal firing of all the neurons with significant on-response (B) and off-response (D) time fields across the first 20 seconds of intertrial delay period. Each row illustrates the normalized firing rate of a single neuron, averaged across all trials, and sorted according to the time of maximum (B) or minimum (D) firing. Plot C shows two example neurons with time fields aligned to the end of the delay period. For each neuron, the start-aligned firing is plotted on the left and the same data re-aligned to the end of the delay is plotted on the right. The neuron in plot 1 had a time field aligned to the start of the delay on the east trials, along with a time field aligned to the end of the delay for both trial types. Plot E shows the percentage of time cells that showed differential firing for the on-response and off-response time cells. Plot F shows the percentage of time cells observed in putative pyramidal neurons and interneurons, with on– and off-responses shown in blue and yellow, respectively.

Interestingly, we found that RSC neurons could also exhibit discrete periods of inhibited firing (Fig. 1A, plots 9-10). These off-responses accounted for 25% (16/65) of the significant time cells seen in the blocked alternation task. These off-response time fields were shorter than the on-responses in terms of their duration (on-responses: 2.95 ± 0.22 sec, off-responses: 1.55 ± 0.12 sec, t(93) = 3.77, p<0.001), but interestingly, they were somewhat more reliable (on-responses: 0.75 ± 0.01, off-responses: 0.97 ± 0.02, t(93) = 7.29, p<0.001) and their contrast between in-field and out-of-field firing was significantly higher (on-responses: 0.38 ± 0.02, off-responses: 0.72 ± 0.04, t(93) = 7.11, p<0.001). However, off-responses were only observed during the first 5 sec of the delay period (Fig.1D) and they were less likely to differentiate the east and west conditions (Fig. 1E). On– and off-responses were not exclusively associated with pyramidal neurons or interneurons. On– and off-responses occurred in similar proportions within the pyramidal neurons and interneurons (Fig. 1F).

Time cells frequently differentiated the ‘go east’ and ‘go west’ blocks of the alternation task. Overall, 80% of time cells (52/65) showed differential firing across the two conditions. Typically, the neurons had a clear time field in one block, while firing in the other block could range from a complete absence of firing during the same epoch in the other block (e.g. Fig. 1A, plots 1-2) to a clear time field, but with reduced firing rate (e.g. plot 6). In some cases, the neuron had time fields in both blocks, but during different epochs (n=9, Fig. 1A, plot 8). Differential firing was more prevalent for the on-response time cells (88%, 43/49) than for the off-response time cells (44%, 7/16).

Time cells were also observed in the cued T-maze task (Fig. 2). However, they were not as prevalent, and they did not differentiate the left and right conditions as frequently as in the blocked alternation task (Fig. 3). Only 15% (31/203) of neurons had significant time cell firing and among these, only seven neurons (23%) differentiated the left and right trials. The overall quality of the time fields also differed across the tasks. Compared to the blocked alternation task, the time fields observed in the cued T-maze were larger (cued T-maze duration = 3.17 ± 0.19 sec, alternation = 2.58 ± 0.17 sec, t(160) = 2.25, p<0.05), less reliable (cued T-maze = 0.67 ± 0.02, alternation = 0.81 ± 0.02, t(160) = 5.64, p<0.001), and had lower in-field to out-of-field contrast (cued T-maze = 0.30 ± 0.02, alternation = 0.47 ± 0.03, t(160) = 4.99, p<0.001). Most of the time cells in the cued T-maze task had on-responses (94% of time cells, 29/31), while only two had off-responses. All seven of the cases of differential firing on left and right trials occurred in neurons with ‘on’ responses. Time fields were seen in pyramidal (n=13) and interneurons (n=18). Only three time cells had a secondary time field and for two of those cells, the secondary time field was present in only one of the trial types while the third one had a secondary time field in both trial types.

**Figure 2.**
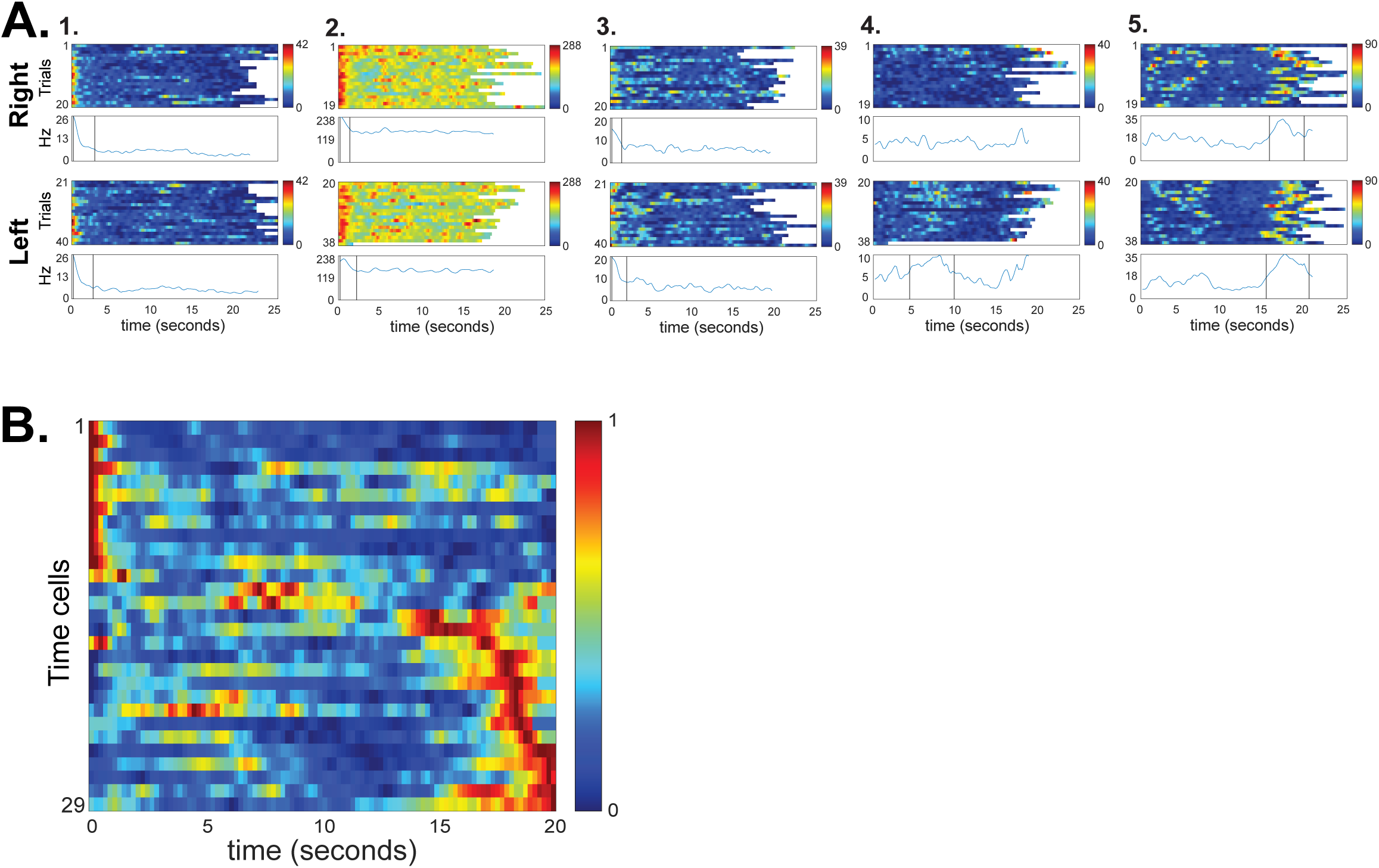
Retrosplenial Time Cells in the cued T-maze task. Plot A illustrates example time cells observed in the cued T-maze task. Firing rate heatmaps are illustrated as in figure 1, with separate plots for right and left trials. Plot B illustrates the temporal firing of all the neurons with significant on-response time fields. Only two neurons were found to have off-response time fields in this task (not shown).

**Figure 3.**
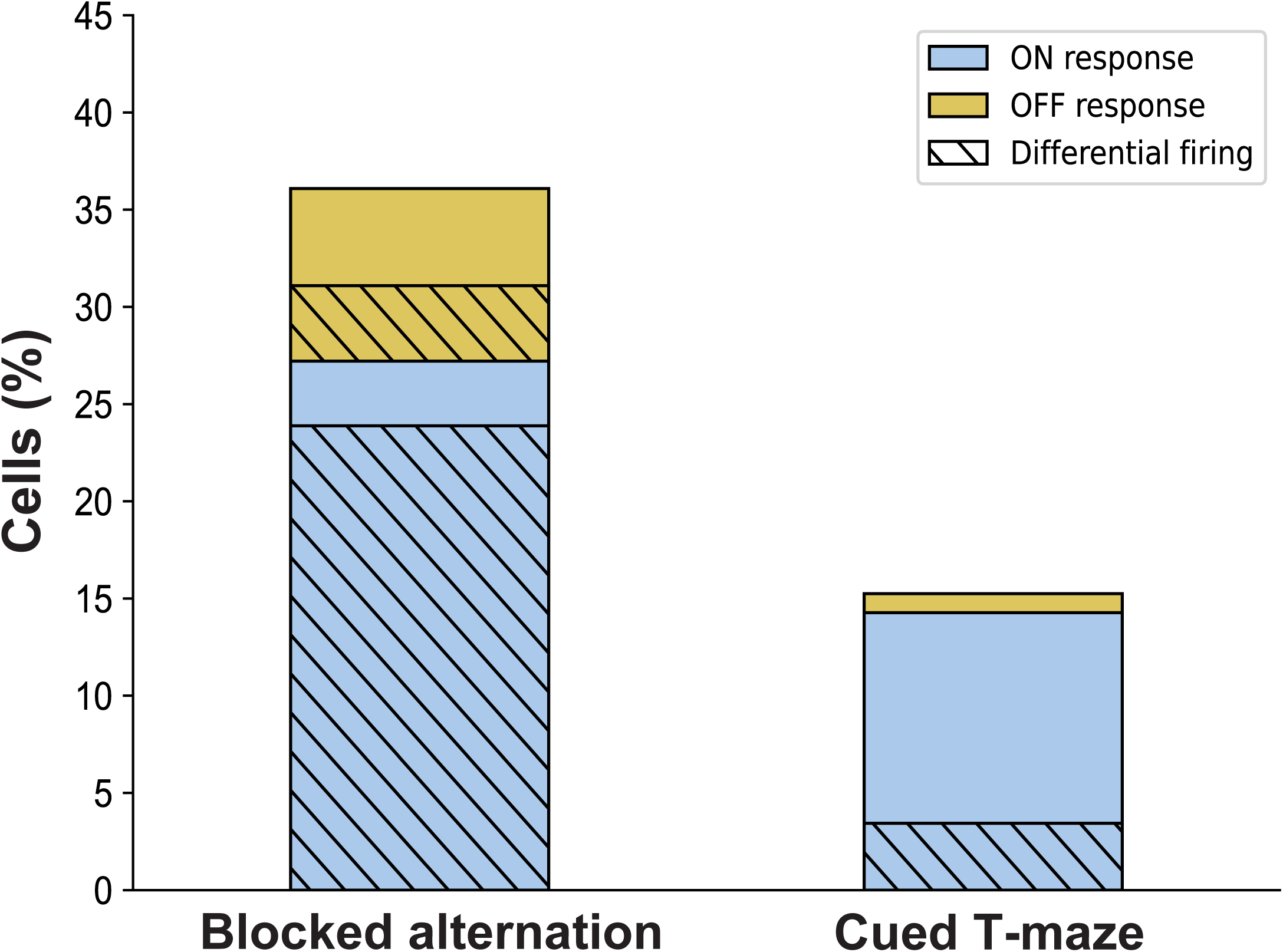
Time cell characteristics differ across behavioral tasks. The percentage of neurons with significant time cells in each task are shown, with on– and off-response time cells shown in blue and yellow, respectively, and differential firing across the east/west or left/right conditions indicated by diagonal lines.

For the above analyses, we compared the delay periods that occurred immediately *after* left and right trials. We reasoned that although there was no requirement that the subject remember the reward locations, firing during the delay period might nevertheless have reflected memory for the events of the previous trial. In contrast, the rats could not anticipate the upcoming trial because the left– and right-rewarded trials were randomized. This feature of the task allowed us to use delay periods that occurred *before* left and right trials as a non-mnemonic control.

When we sorted left and right trials that way, we still found that 15% (30/203) of the neurons had time fields and 20% (6/30) of them differentiated the left and right conditions. The similar number of time cells suggests that even the relatively small percentage of time cells seen in our original analysis (above) are not related to memory for the trials.

RSC neurons are known to exhibit spatial firing (Alexander & Nitz, 2015; Miller et al., 2019) and many of the time cells observed here also showed spatial firing during the trials (Fig. 4). As in previous studies (e.g. Gill et al, 2011), time cell firing did not appear to be related to spatial firing during the trials. We examined this by binning the firing rates into 40 spatial bins along the trajectory for each trial from the start to the reward and correlating the firing rates with the time binned firing rates used to assess time fields. As expected, the spatial and temporal firing rates were not correlated (mean Pearson’s r = 0.0007 ± 0.005), suggesting that temporal firing is not a recapitulation of spatial firing on the maze.

**Figure 4.**
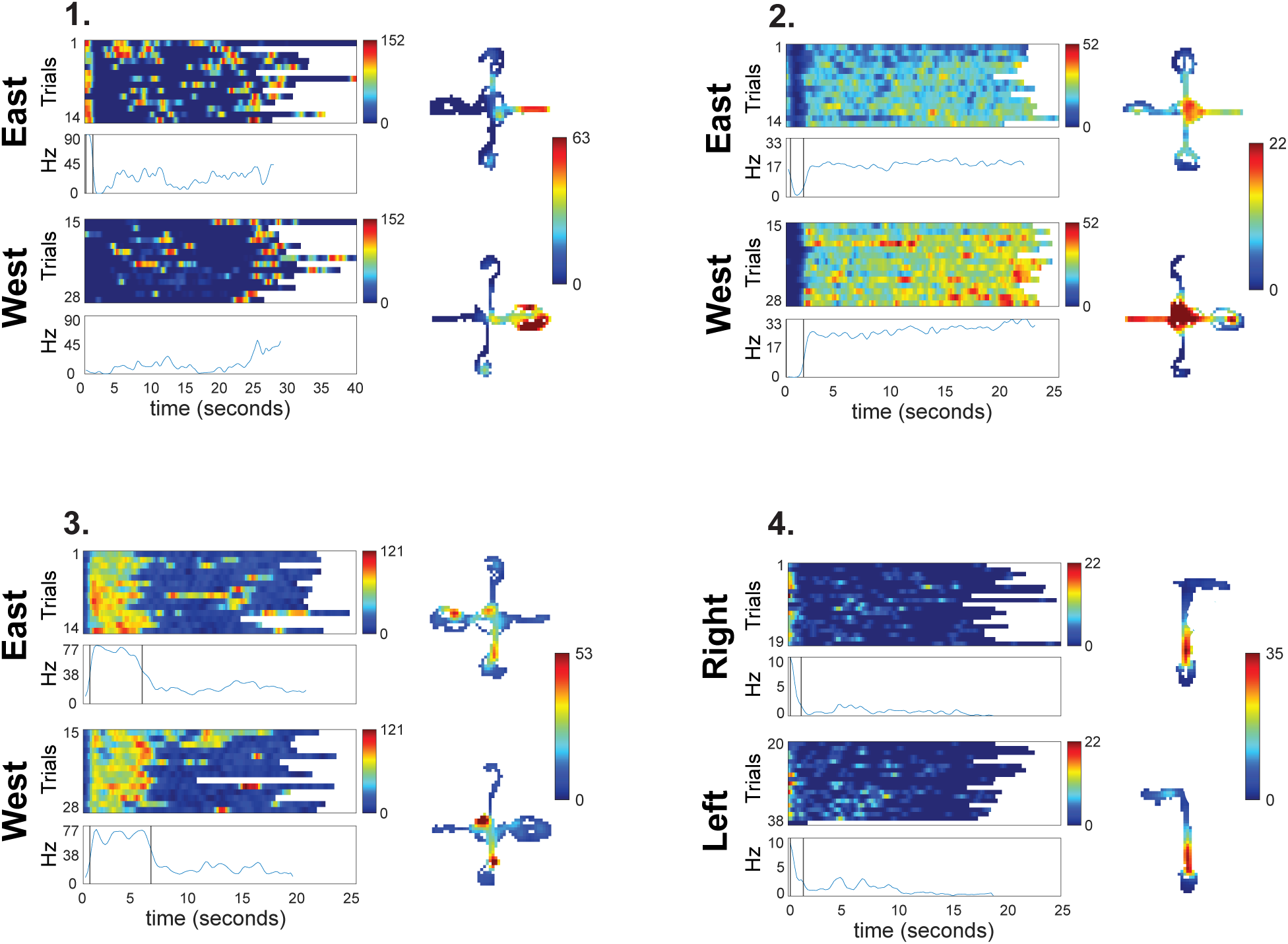
Spatial firing in RSC time cells. Spatial firing patterns observed during the trials are shown for four example neurons with time fields during the intertrial delay period. Plot 1 is the same neuron illustrated in figure 1C, plot 1. Plot 4 is an example neuron from the cued T-maze task.

## Discussion

Time cells were a prevalent feature of RSC representations, and their time fields were similar to those previously observed in the hippocampus (Gill et al., 2011; MacDonald et al., 2011; Pastalkova et al., 2008) and the medial entorhinal cortex (Heys & Dombeck, 2018; Kraus et al., 2015). These responses took the form of discrete periods of reliably elevated, or suppressed, firing with high contrast to the background firing rate. The on-response time fields covered the entire intertrial delay period and they differentiated the east and west conditions of the blocked alternation task, which is also similar to hippocampal time fields (Gill et al., 2011). Interestingly, more than one third of RSC neurons had time fields in the blocked alternation task, compared to about 20% of hippocampal neurons in the same task (Gill et al., 2011), suggesting that time cells may be more prevalent in the RSC.

Off-response time fields were unexpected, as they have not been reported before. These responses are possible in the RSC because, unlike the hippocampus, RSC neurons often have high baseline firing rates which allows responses to take the form of either increases or decreases from the baseline. Previous studies of the RSC have found reliable off-responses to critical stimuli and events (Smith et al., 2012; Vedder et al., 2017) and these responses likely contribute importantly to population level representations of space and context (Miller et al., 2019, 2021). The off-response time fields were discrete and reliable, and they differentiated the east and west conditions of the blocked alternation task at a rate similar to our previous study of hippocampal time cells in this task (Gill et al., 2011). However, there were some potentially important differences between the on– and off-response time cells. Differential firing was far less common in the off-response neurons (Fig. 3) and off-response time fields were only seen during the first five seconds of the delay period (Fig. 1D). However, the relative rarity of these responses in the population makes it difficult to determine whether longer latency time fields are completely absent or simply less common than early time fields.

The task differences observed here are consistent with the idea that time cells are sensitive to the memory demands of the task. Time cell firing was both more prevalent and more likely to differentiate the east and west conditions in the blocked alternation task, as compared to the cued T-maze task. The blocked alternation task has a clear memory requirement insofar as the rats must hold two distinct goal memories in preparation for the ‘go east’ and ‘go west’ trials. In contrast, the cued T-maze task simply requires the rat to approach the light cue for reward, a single procedural memory which likely does not engage the episodic memory system (Lee et al., 2008; McDonald & White, 1994). Consistent with this idea, our previous study (Vedder et al., 2017) found that RSC neurons respond to the light cue the same way, regardless of whether it was presented on the left or the right. Similarly, we found that differential time fields were rare in this task.

The task differences we observed are notably similar to previous reports on the hippocampus, where time cells were found in tasks that require subjects to hold different memories during the delay period, but not in similar tasks which do not (Gill et al., 2011; Pastalkova et al., 2008; but see Sabariego et al., 2019). Interestingly, 80% of the RSC time cells differentiated the east and west trials, as compared to about half of hippocampal time cells recorded in the same task (Gill et al., 2011), suggesting that memory related time cell coding might be more prevalent in the RSC.

Our observation of time cell coding is consistent with an RSC role in episodic memory (Ranganath et al., 2005; Steinvorth et al., 2006). Previous authors have suggested that hippocampal time cells play a role in encoding the temporal context, which provides a mechanism for encoding the sequence of stimuli and events that define an episodic memory (Eichenbaum, 2014; Howard & Eichenbaum, 2013). Our results suggest that the RSC may play a key role in this process. This is consistent with findings from patient T.R. whose left-RSC lesion produced a remarkable impairment in temporal memory (Bowers et al., 1988). The deficit was a specific failure to remember which items came earlier or later within lists of to be remembered items, even though recognition of the individual items was unimpaired. Bowers and colleagues (1988) describe this impairment as a failure of time-tagging, which echoes the idea of temporal context as a mechanism for sequencing items in memory (Howard & Kahana, 2002). In a notably similar pattern of results, rats with RSC lesions were found to be impaired in memory for the temporal order of objects, but not for recognition of the objects themselves (Powell et al., 2017). Relatedly, damage to the anterior thalamic nuclei, a primary input to the RSC, in Korsakoff’s amnesia also produces deficits in temporal memory (Hunkin & Parkin, 1993; Kopelman, 2015; Kopelman et al., 1997; also see Nelson, 2021). Finally, studies of instrumental discrimination learning have shown that RSC neurons exhibit systematic time-dependent changes in stimulus-evoked responses, which are thought to underlie a similar kind of temporal context (Freeman & Gabriel, 1999).

The finding of time cell coding in the RSC provides an additional point of functional similarity between the RSC and the hippocampus, along with spatial and contextual memory, which highlights the importance of understanding the roles of these structures as components of a coherent memory circuit (Smith et al., 2022). The source of RSC time cell information is not known, but hippocampal input is known to be critical for some kinds of RSC representations (Cowansage et al., 2014; De Sousa et al., 2019; Katche et al., 2013; Mao et al., 2018; Milczarek et al., 2018) and there is evidence that hippocampal input influences RSC activity (Alexander et al., 2018, Kang & Gabriel, 1998). Moreover, findings of time cells in the entorhinal cortex, prefrontal cortex and the striatum (Akhlaghpour et al., 2016; Heys & Dombeck, 2018; Kraus et al., 2015; Mello et al., 2015; Ning et al., 2022; Tiganj et al., 2017) suggest that, whatever the source of this signal, time coding is widespread in brain regions known to be involved in learning and memory.

## Acknowledgements

This work was supported by MH083809 to D. Smith.

## Notes

### Competing Interest Statement

The authors have declared no competing interest.

